## Supplementary Figure for "Exploring cellular changes in ruptured human quadriceps tendons at single-cell resolution"

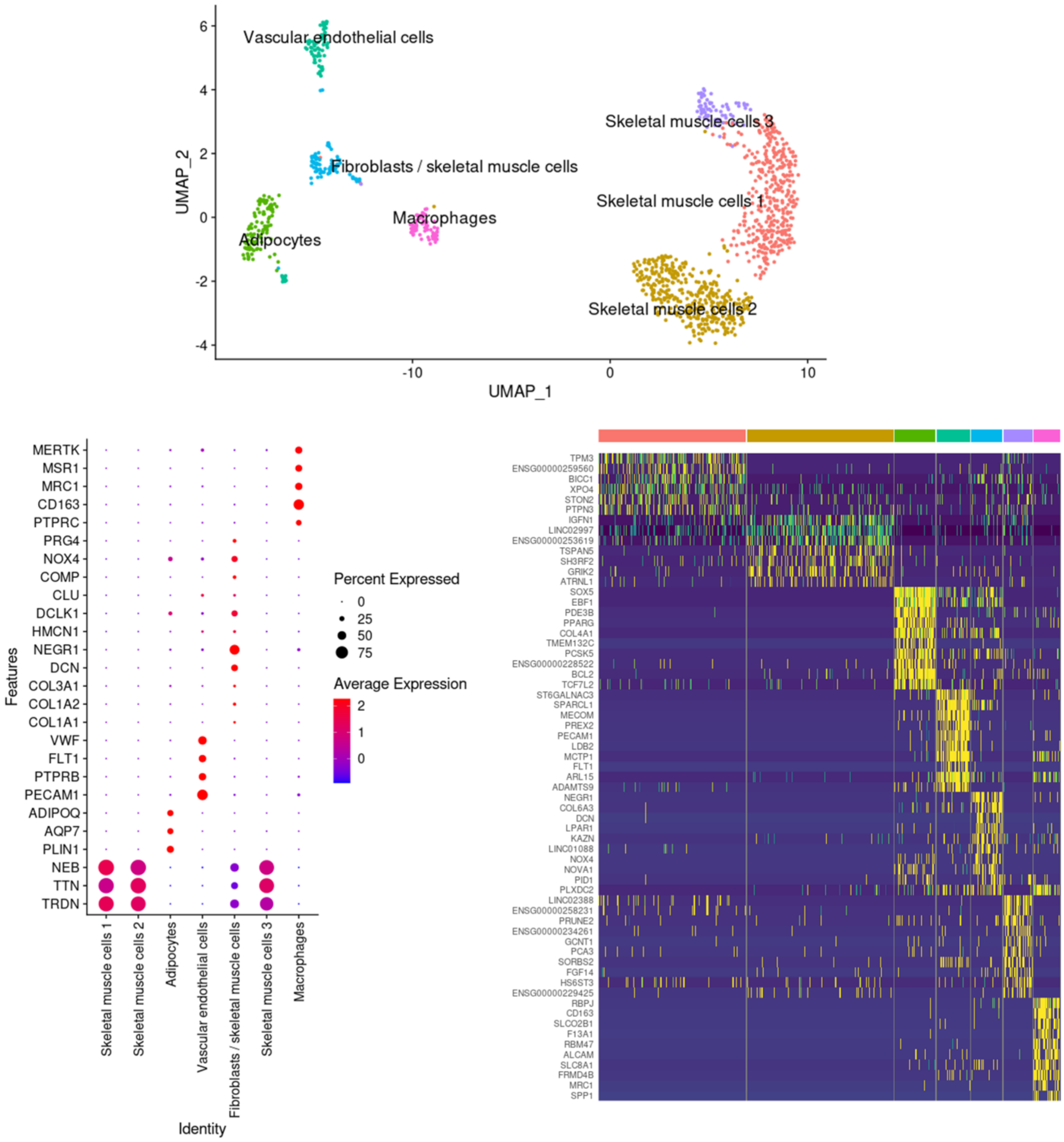


**Supplementary Figure 1.** Cell clusters and gene expression of canonical markers in data from patient MSK1250.

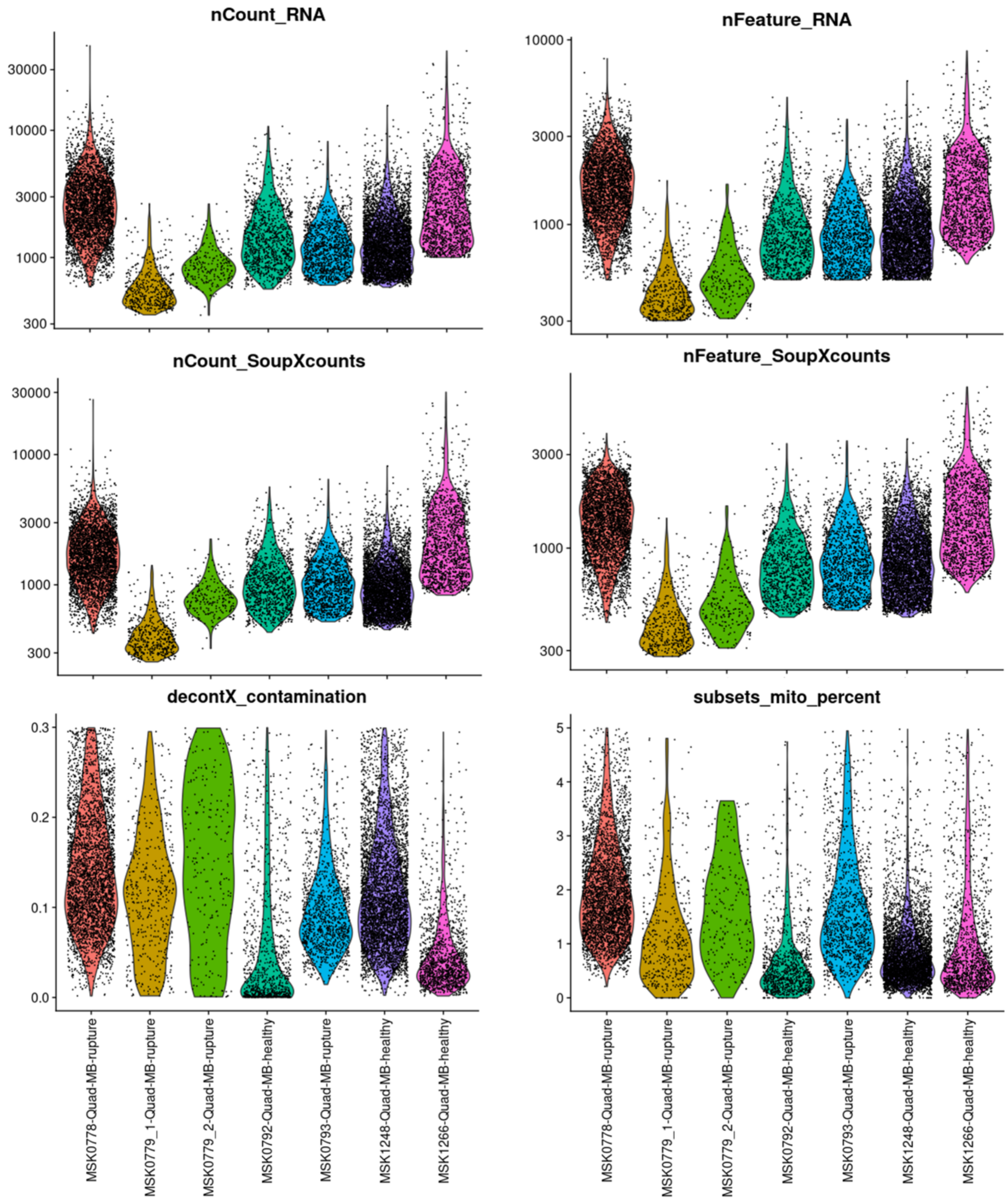


**Supplementary Figure 2**. QC metrics of each dataset, including number of counts (nCount) and number of features (nFeature) in the RNA and SoupXcounts assays, decontXcontamination score, and percentage of mitochondrial reads (subsets_mito_percent).

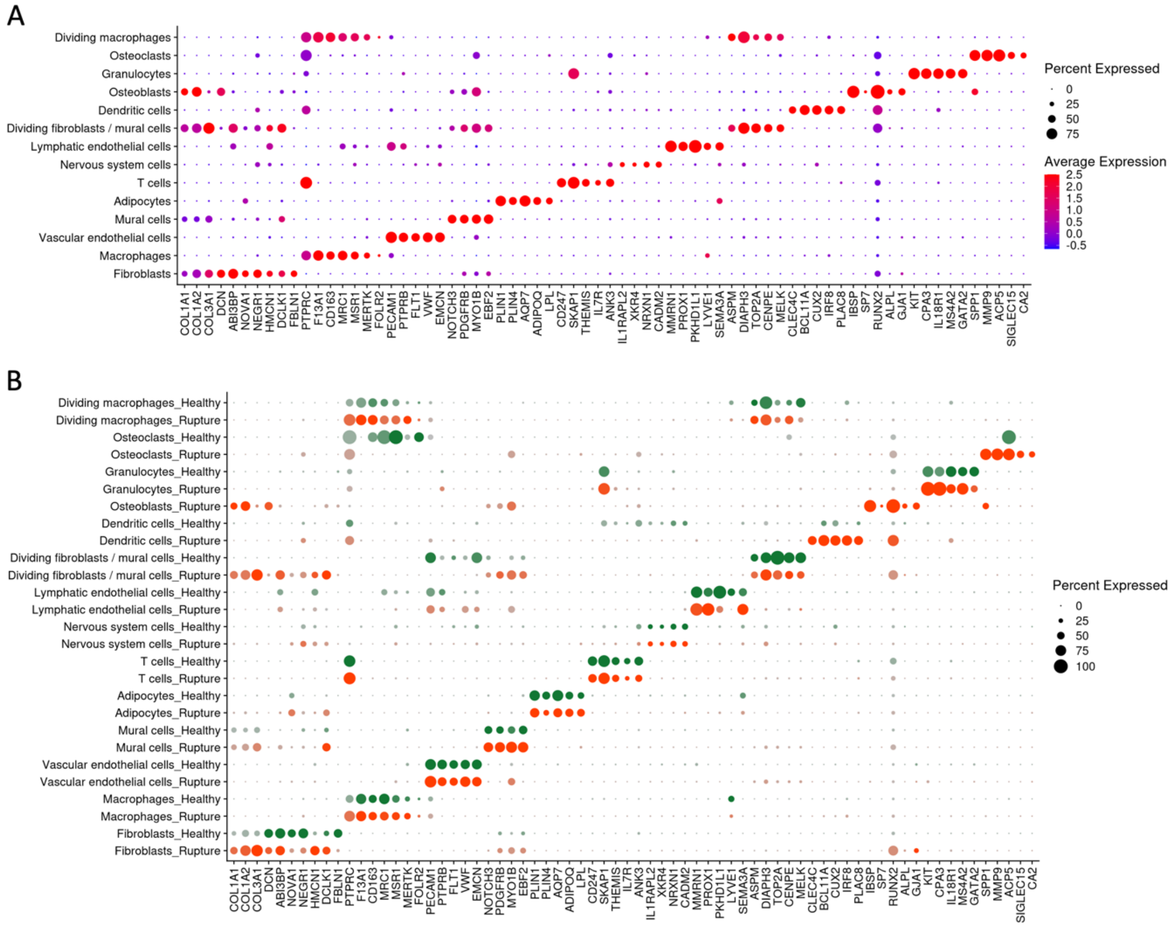


**Supplementary Figure 3.** Expression of canonical markers in each identified cell subset.

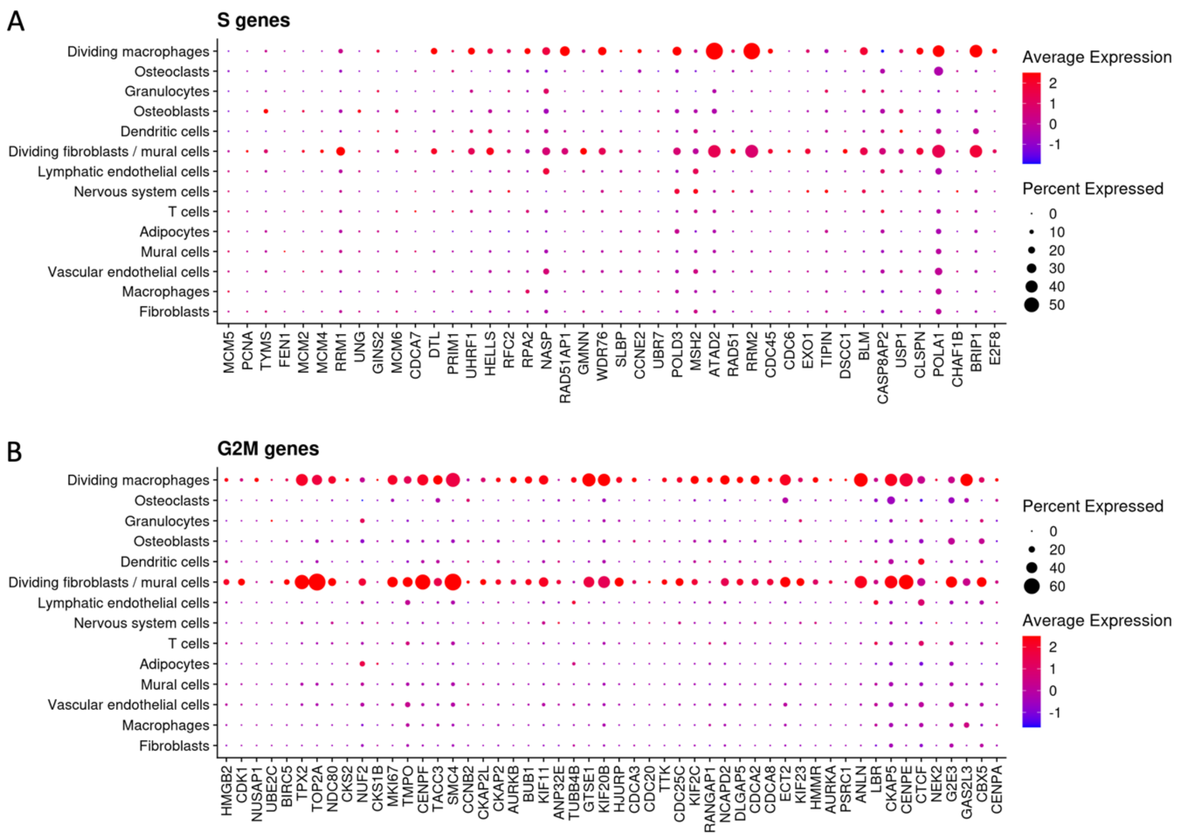


**Supplementary Figure 4**. Expression of S- and G2M-phase genes in each cell subset as presented in Figure 1.

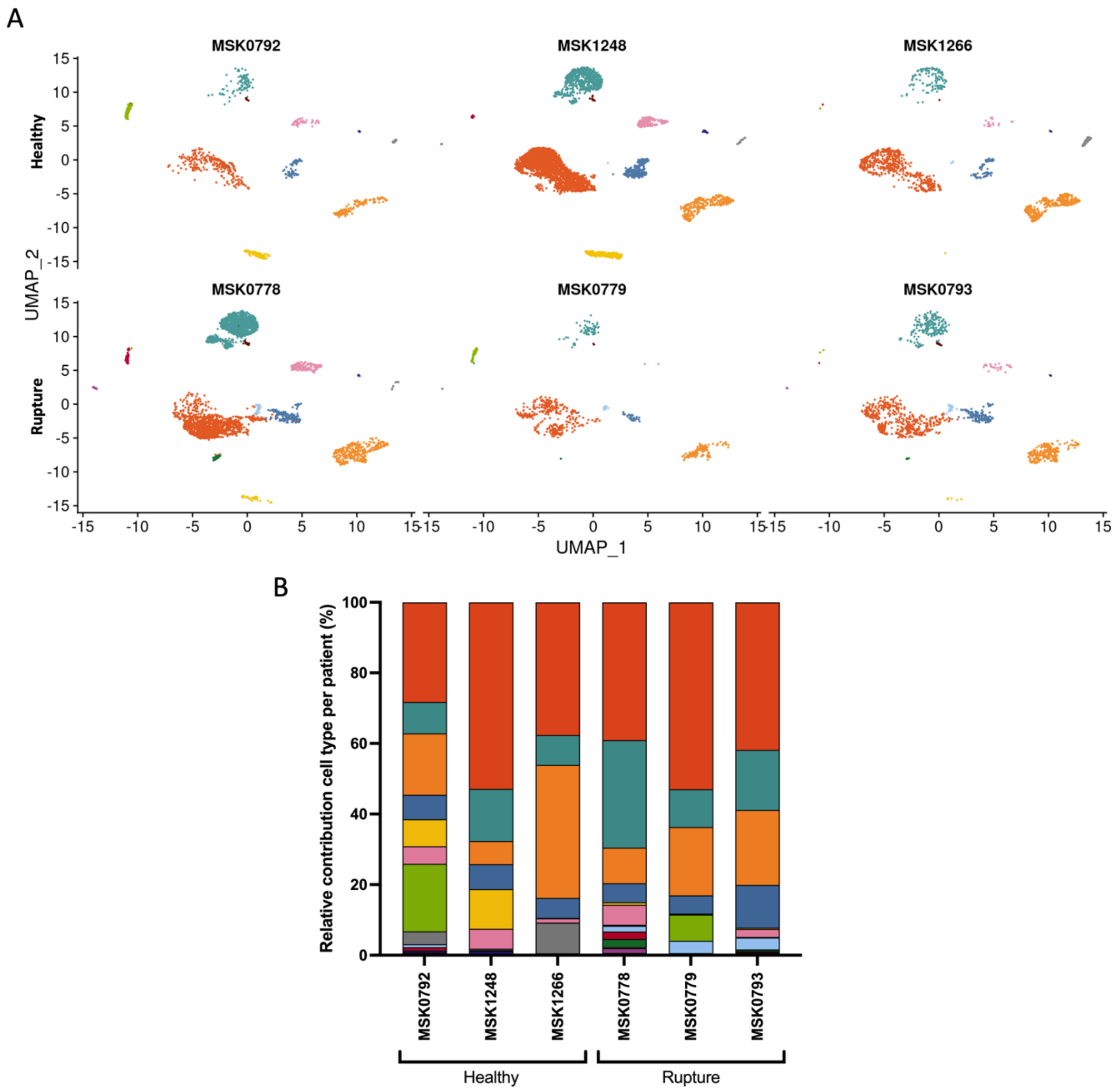


**Supplementary Figure 5**. Single nucleus RNA sequencing results by disease status and patient. (A) UMAP embedding of 12,817 nuclei from quadriceps tendons split by patient; top row are healthy donors and bottom row are donors of ruptured quadriceps samples. (B) Mean cell type frequency per patient.

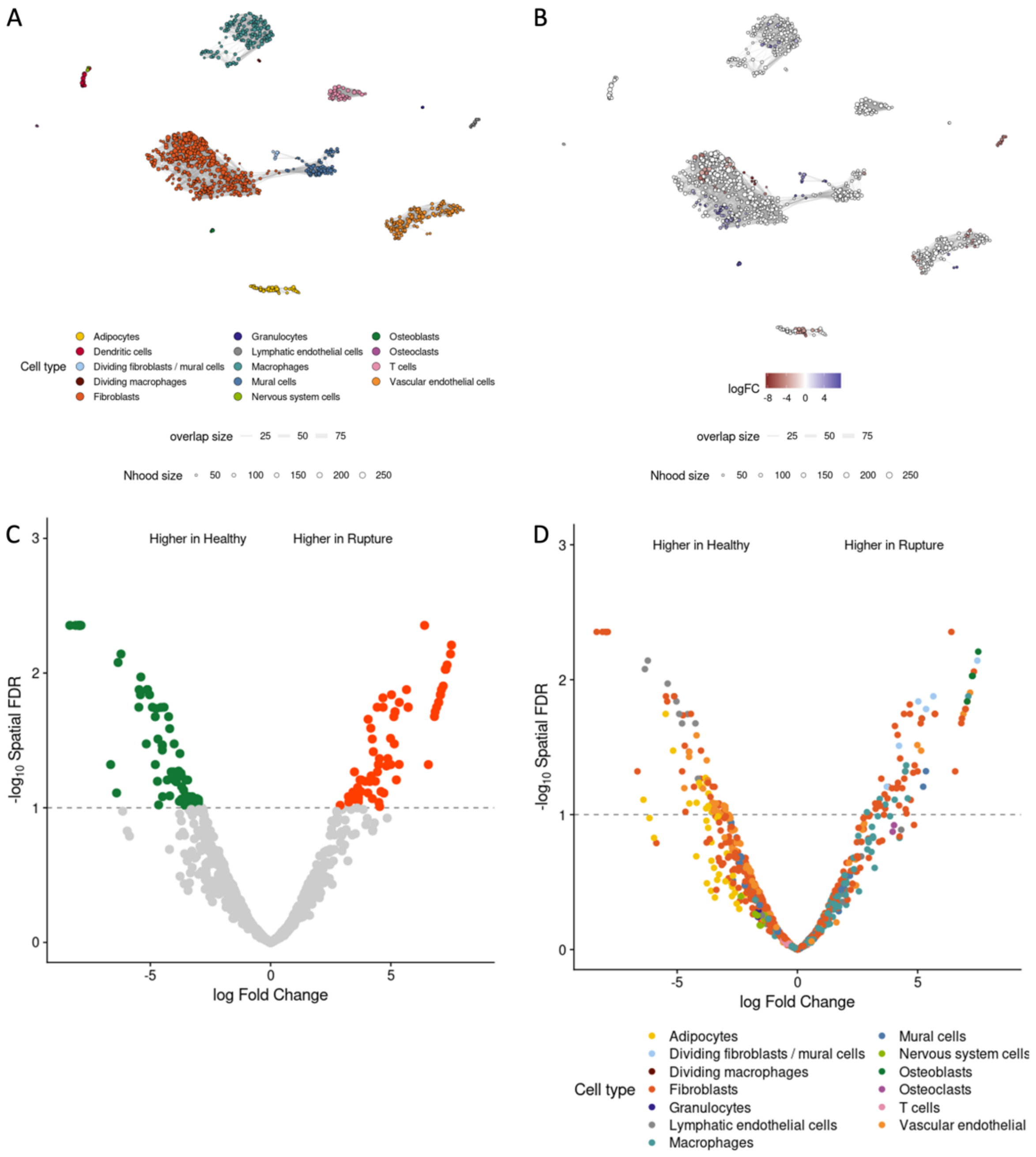


**Supplementary Figure 6**. MiloR differential abundance analysis. MiloR was used to assign neighbourhoods (A) and test these neighbourhoods for differential abundance (B). (C-D) Volcano plots depict the differential abundance of each neighbourhood, either coloured by (C) tendon disease (ruptures (red) or healthy (green)), or by (D) cell type.

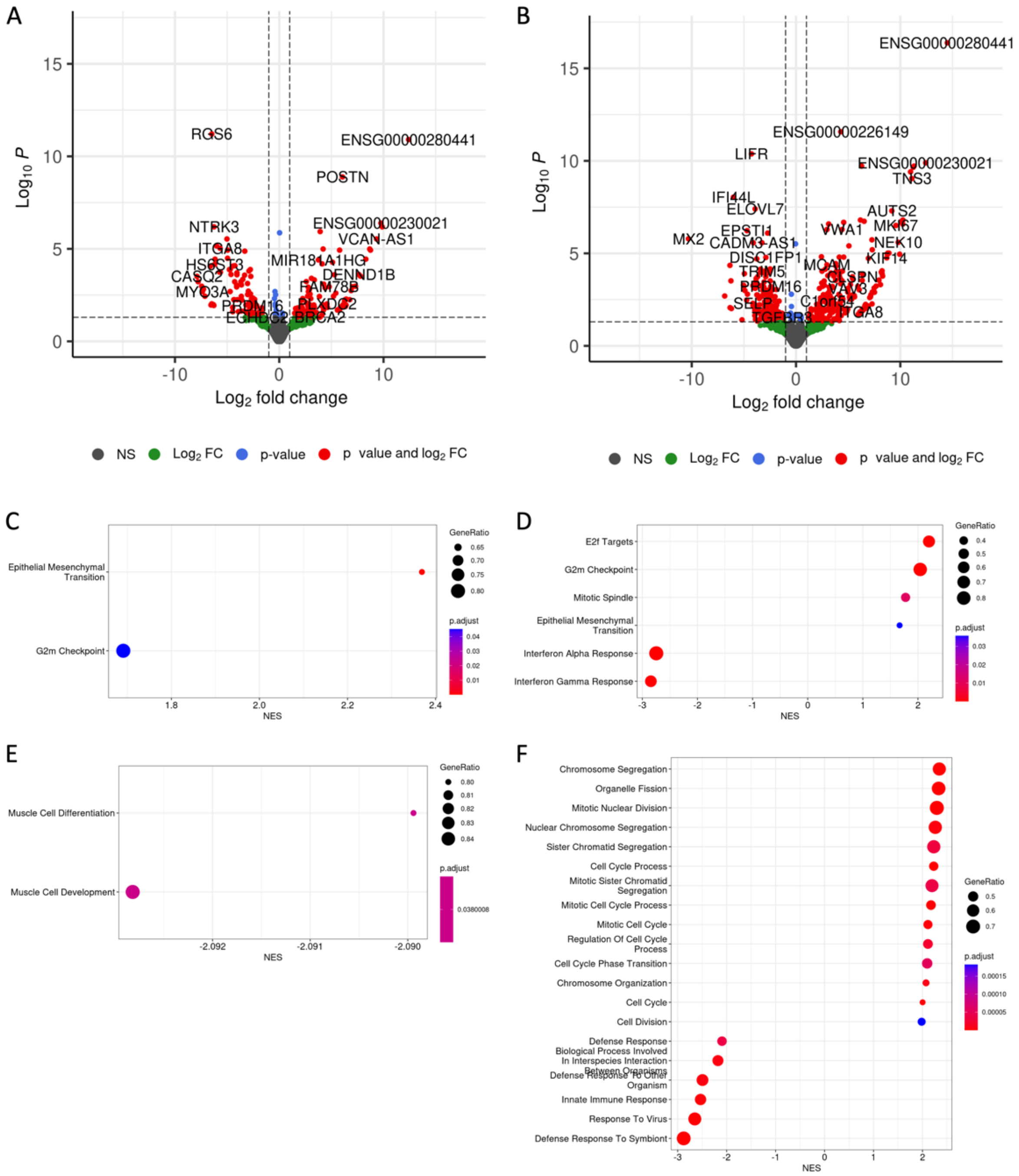

**Supplementary Figure 7**. Differential gene expression results of mural cells (left) and vascular endothelial cells (right), including volcanoplots (A-B), Hallmark pathway results (C-D), and Gene Ontology Biological Processes results (E-F), either all (E) or top 20 (F).

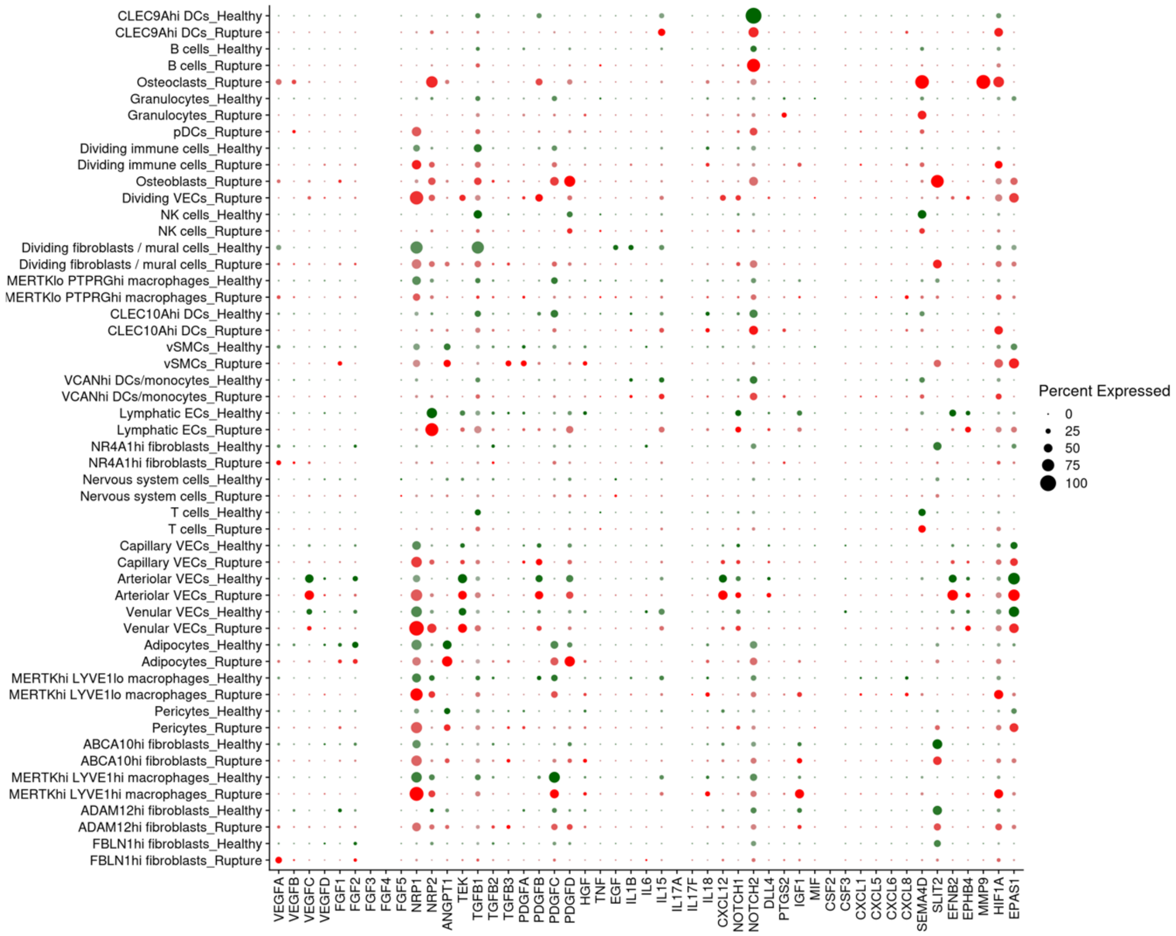


**Supplementary Figure 8**. The expression of pro-angiogenic factors by all the identified clusters, split by healthy (green) and rupture (red).

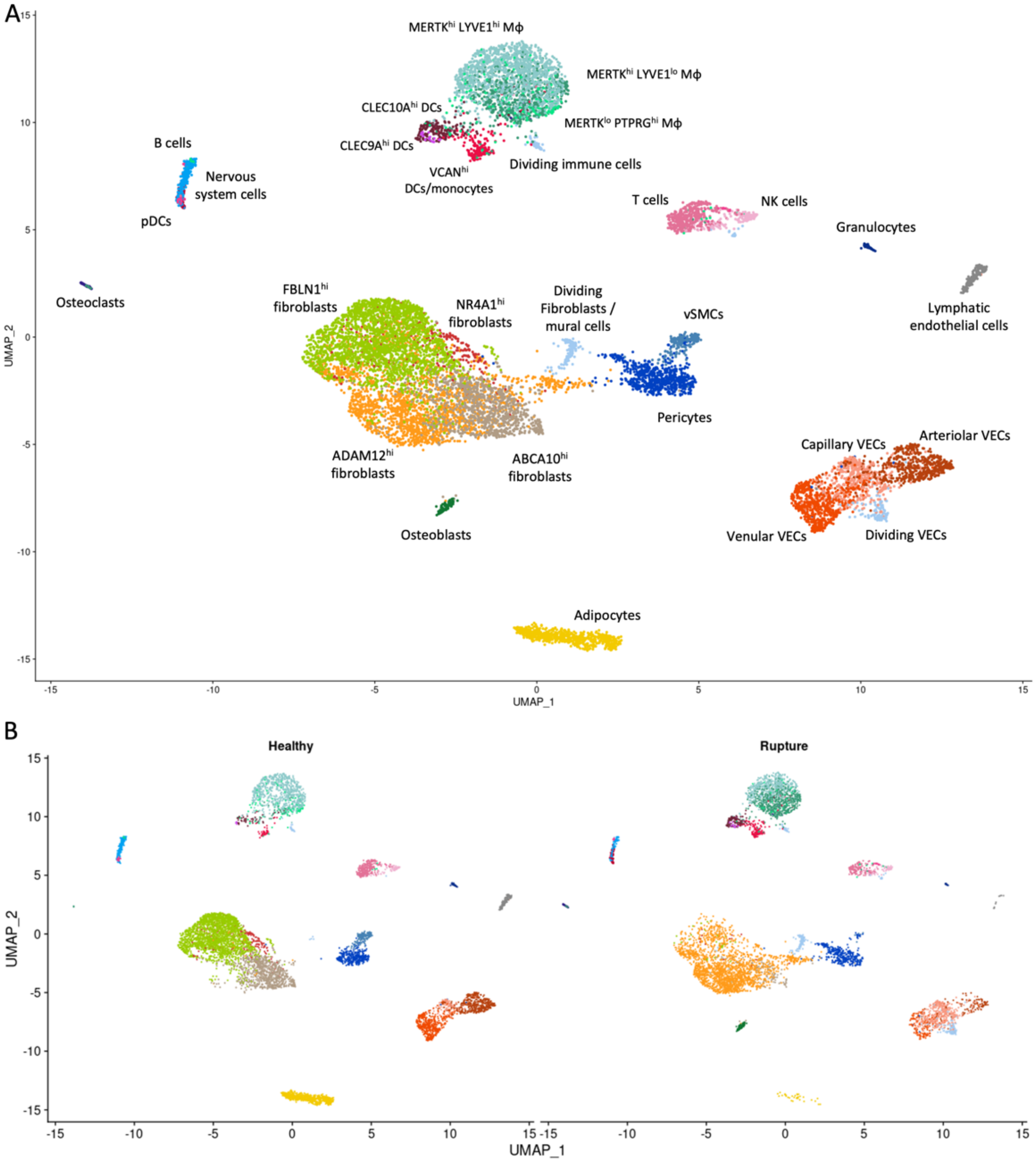


**Supplementary Figure 9**. UMAP embedding with updated cell annotation based on the subclustered fibroblast, endothelial, and immune cell types, either overall (A) or split by tendon disease (B). DC = dendritic cells, MΦ = macrophages, VEC = vascular endothelial cells, vSMCs = vascular smooth muscle cells.

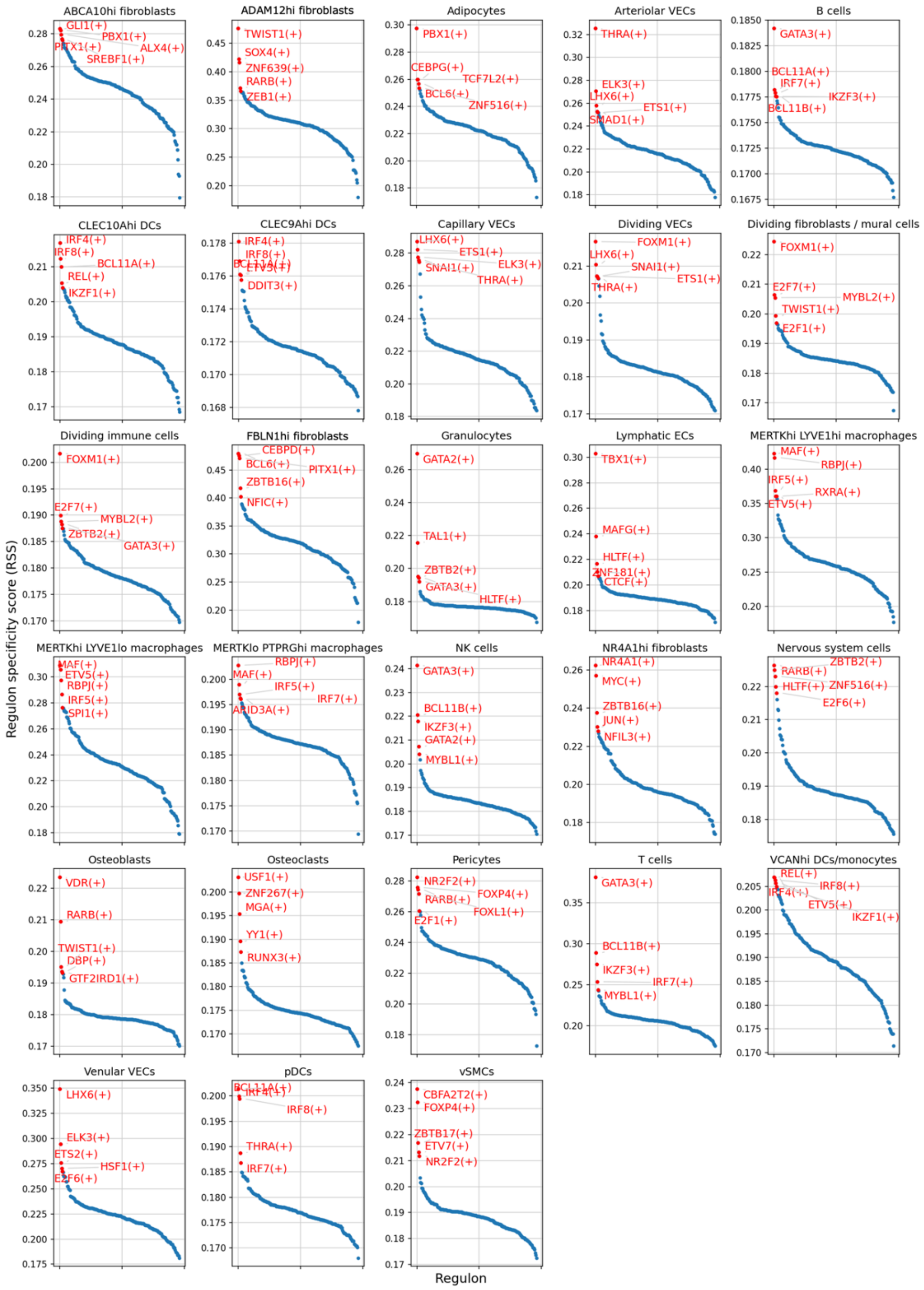


**Supplementary Figure 10**. Regulon specificity scores (RSS) for each of the identified cluster. For each cluster, the top 5 regulons are highlighted in red.
